# Oxytocin modulates behavioral economic and dopaminergic measures of motivation to seek intravenous oxycodone in rats

**DOI:** 10.64898/2026.10.16.759686

**Authors:** Harrison L. Blount, Madison P. Gits, Elizabeth A. Colley, Jennifer E. Kang, Courtney S. Wilkinson, Marek Schwendt, Lori A. Knackstedt

**Author notes:** **Corresponding author:**Lori A. Knackstedt, PhD, Department of Psychology, University of Florida, Gainesville, FL 32611, USA.

## Abstract

The opioid epidemic persists despite available pharmacotherapies for opioid use disorder (OUD), motivating interest in treatments targeting the motivational and neurochemical substrates of opioid seeking. Peripherally administered oxytocin (OT) attenuates drug seeking and consumption across substances and modulates the mesolimbic dopamine system, yet its effects on opioid seeking and its underlying neurobiology remain poorly characterized. Here we investigated the effects of OT on behavioral and dopaminergic measures of opioid reinforcement using *in vivo* fiber photometry with the dopamine sensor, GRAB_DA2m_, in the nucleus accumbens (NA) core of male and female rats. Daily OT (15 IU/kg, IP) administered 30 min prior to operant sessions increased demand elasticity for intravenous oxycodone self-administration in males but not females. Acute OT (15 and 45 IU/kg, IP) increased GRAB_DA2m_ dopamine signal in the NA core in females, and potentiated oxycodone-evoked dopamine release in both sexes. In a high-effort task, NA core dopamine increased prior to lever presses, and acute OT (15 IU/kg, IP) produced sex-divergent effects on phasic NA core dopamine release during lever pressing, increasing peak signal and area under the curve measures in males. Support vector machine classifiers determined that features of dopamine transients (e.g., mean peak, standard deviation of the amplitude, peak-valley range) before an active lever press distinguished between treatment and sex. Thus, the ability of chronic oxytocin to reduce the motivation to self-administer oxycodone in male rats is independent of oxycodone-induced NA dopamine release but rather may involve dopamine transmission during lever pressing under high effort conditions.

## Introduction

Opioid use disorder (OUD) remains prevalent in the United States with great economic, and health burdens (*1*). Although existing FDA-approved treatments are effective, their impact is limited by accessibility, persistent craving, and high rates of attrition (*2–5*). Identifying well-tolerated pharmacotherapies that reduce opioid use remains a public health priority.

Oxytocin (OT) is a neurohormone with emerging therapeutic potential for substance use disorders, including opioid use disorder (*6*, *7*). Acutely administered OT consistently reduces consumption of natural and drug rewards (e.g., food, sucrose, alcohol, cocaine, heroin) across animal models (*8–14*). The results of clinical trials using OT interventions for OUD are more heterogeneous, but decreased opioid craving is reported following oxytocin administration in two of five existing studies (*15–19*). While acute oxytocin reduces cued heroin seeking (*20–22*), whether chronic OT treatment can alter ongoing motivation to seek/take opioids remains unknown. Thus, here we sought to assess the ability of OT to reduce the motivation to seek intravenous oxycodone, one of the most widely abused prescription opioids (*23*, *24*). A behavioral economic assessment of the effects of OT on the motivation to seek oxycodone was conducted, an approach can be used across species, allowing demand curve parameters such as elasticity of demand (alpha or “α”) to serve as measures of drug seeking that are sensitive to pharmacological and neurobiological manipulations (*25–30*).

Chronic opioid exposure induces neuroadaptations within mesocorticolimbic circuitry that strengthen drug-cue associations and promote addictive behaviors (*31–33*). The nucleus accumbens (NA) is a critical component of the mesocorticolimbic circuitry mediating opioid reward, self-administration, and relapse-like behaviors (*34–38*). Intra-NA core inhibition of D1- or D2-like dopamine receptors reduces cue-induced heroin seeking (*37*, *38*), and lesions of NA core dopamine-terminals attenuate heroin self-administration under second-order schedules of reinforcement (*36*). Thus, targeting dopamine transmission in the NA core represents one potential method by which to reduce the motivation to seek opioids.

OT receptors are expressed throughout the nervous system, including on ventral tegmental area (VTA) dopamine neurons and their activation potentiates downstream dopamine release (*39*– *44*). OT also alters dopamine release in the context of drugs of abuse; acute OT prevents alcohol from increasing NA shell dopamine efflux in male rats, and in cocaine-experienced male rats, OT increases NA core DA (*45–47*). However, these studies utilized microdialysis procedures, which quantify neurotransmitter levels at the minute-time scale. Genetically encoded dopamine sensors such as GRAB_DA_ offer ligand sensitivity and rapid kinetics needed to resolve neurotransmission on the millisecond timescale. Coupled with operant responding in freely moving subjects, *in vivo* fiber photometry is well suited to characterize neurotransmitter dynamics during drug seeking (*48*, *49*).

Here we first tested the hypothesis that OT affects motivation for intravenous oxycodone, finding that chronic OT increased demand elasticity for oxycodone in males but not females. Phasic NA core dopamine release encodes reward prediction and motivational salience and facilitates the vigor of operant responding (*50–53*), and acute oxycodone increases NA core dopamine (*47*). Thus, we used *in vivo* fiber photometry with the GRAB_DA2m_ sensor to test the hypothesis that NA core dopamine responses to intravenous oxycodone and lever pressing for oxycodone would be differentially affected by OT in males and females.

## Materials and Methods

### Animals

Male and female Sprague Dawley rats (N = 70; Charles River Laboratories; 8 weeks old at arrival) were used in these experiments. Rats were housed in climate-controlled cages under a reverse 12-h light/dark cycle. Rats were food-restricted (16-20 g/day) to promote operant learning with ad libitum water. All procedures were performed during the dark phase, complied with National Institutes of Health guidelines, and were approved by the University of Florida Institutional Animal Care and Use Committee.

### Drugs

Oxycodone HCl was obtained via the NIDA Drug Supply Program (Research Triangle Institute) and was dissolved in 0.9% physiological saline. The vehicle (VEH) for OT (15 IU/mg, Sigma Aldrich: O4375) was 0.9% physiological saline. OT was administered at doses of 15 and 45 IU/kg, equivalent to 1 mg/kg and 3 mg/kg (*9*, *14*, *55*). IU/kg dose terminology was utilized herein to align with clinical and cross-species OT dose reporting practices (*56*, *57*).

### Surgeries

Jugular vein catheter implantation and post-operative care were performed as previously described (*47*). In photometry experiments, 1 *μ*L of AAV9-hsyn-GRAB_DA2m (titer 1 × 10^12^ vg/mL; Addgene; #140553-AAV9) was infused unilaterally into the NA core (female: AP +1.5, ML ±1.5, DV −5.8 mm; male: AP +1.5, ML ±1.8, DV −6.7 mm from bregma), see Fig. 2B (*58*) at an infusion rate of 0.14 *μ*L/min. The microinjector remained in place for 8 minutes (min) following infusion. An optical cannula (400 *μ*m, 0.66 NA, ceramic, Doric Lenses) was then implanted over the AAV site and secured to the skull using screws and dental cement (*47*). Rats recovered for three weeks prior to photometric recordings and, during this time, were habituated to patch cord tethering.

### Experiment 1: Chronic oxytocin and oxycodone economic demand

Rats (female n = 16, male n = 16) were trained to intravenously self-administer (IVSA) oxycodone for 3 hours/day and 6 days/week in two-lever operant chambers (Med Associates) under fixed-ratio (FR) schedules of reinforcement, with six days of responding on an FR1 followed by six days of the FR3 schedule. Active lever presses delivered a 0.1 mg/kg oxycodone infusion paired with audiovisual cues (5-second stimulus light, 2900-Hz tone) and were followed by a 20- s timeout during which time active lever presses did not deliver drug or cues. Inactive lever presses were recorded but had no programmed response.

After training, chronic OT’s effect on economic demand for intravenous oxycodone was evaluated (see Fig. 1A). The dose of OT (15 IU/kg; also referred to as 1 mg/kg in many studies) was selected based on its ability to acutely decrease alcohol intake and cued heroin seeking (*9*, *20*). FR requirements were increased on a quarter- and eighth log scale every second session until rats failed to obtain a single infusion (*54*). OT/VEH was administered intraperitoneally (IP) 30 min before IVSA sessions, as sedative effects of OT diminish 30 min after administration (*14*, *45*, *59*, *60*). Daily OT/VEH injections began at FR24 for males, as a drop in intake was hypothesized to occur immediately. Upon finding a delayed effect in males, OT administration began at FR10 for females. Rats were assigned to OT/VEH groups such that training intake did not differ between conditions.

**Fig. 1.**
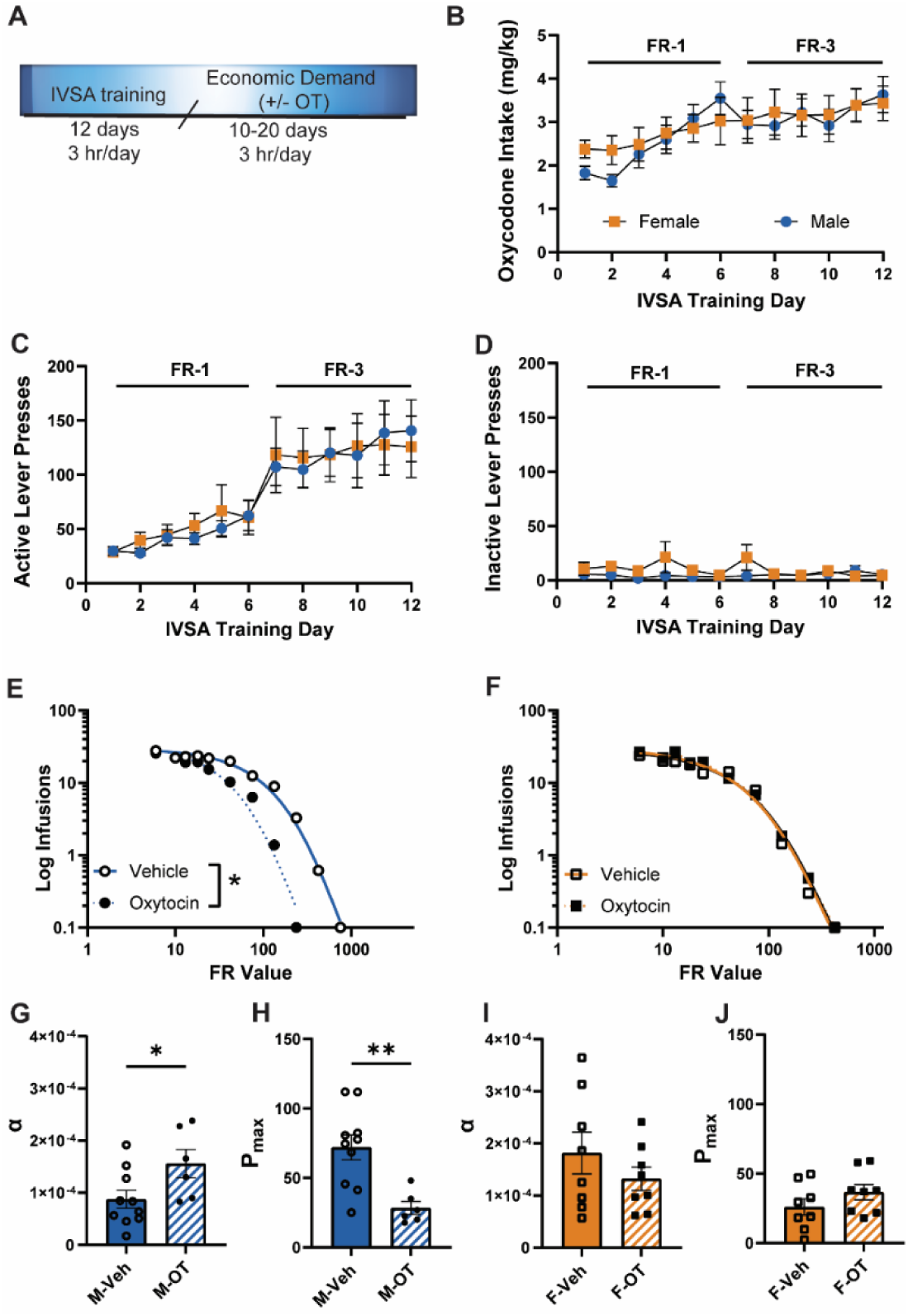
Chronic oxytocin increases demand elasticity for intravenous oxycodone in male rats. **A.** Experimental timeline. Oxycodone intake **(B)**, active lever presses **(C)**, and inactive lever presses **(D)** did not differ between females and males during self-administration training. Daily 15 IU/kg oxytocin (OT) increased demand elasticity for intravenous oxycodone in male rats **(E)** but not female rats **(F)**. Alpha and P_max_ were computed from demand curves. **G**. Alpha (α) was increased by chronic OT in males, indicating more elastic demand for oxycodone. **H**. The price at which consumption becomes elastic, P_max_, was decreased by OT in males. Alpha was not altered by OT in females (**I**), nor was P_max_ (**J**). * = p < 0.05; ** = p < 0.01 VEH v. OT.

### Experiment 2: In vivo fiber photometry assessment of dopamine dynamics following acute oxytocin and non-contingent oxycodone

A separate cohort of rats (female n = 11, male n = 12) with jugular catheters and NA core GRAB_DA2m_ were habituated to patch cord tethering in custom plexiglass experimental chambers (16×16×15 inch). Using a within-subjects design, rats underwent three photometry recordings following 0, 15, or 45 IU/kg OT (IP), administered in a randomized fashion and on alternating days. A higher dose (45 IU/kg; also referred to as 3 mg/kg in many publications) was also used here, as the 15 IU/kg/day dose was ineffective in altering motivation for oxycodone in females in Experiment 1, and 45 IU/kg attenuates cocaine self-administration (*10*, *59*). During each recording session, rats were habituated to the chamber for 5 min, followed by a 70-minute recording session. After a 10-minute baseline recording, rats received either OT or VEH and 30 min later, a dose of noncontingent oxycodone (0.5 mg/kg, IV) that increases GRAB_DA2m_ signal for 30 min after infusion (*47*). See timeline of recording sessions in Fig. 2A.

**Fig. 2.**
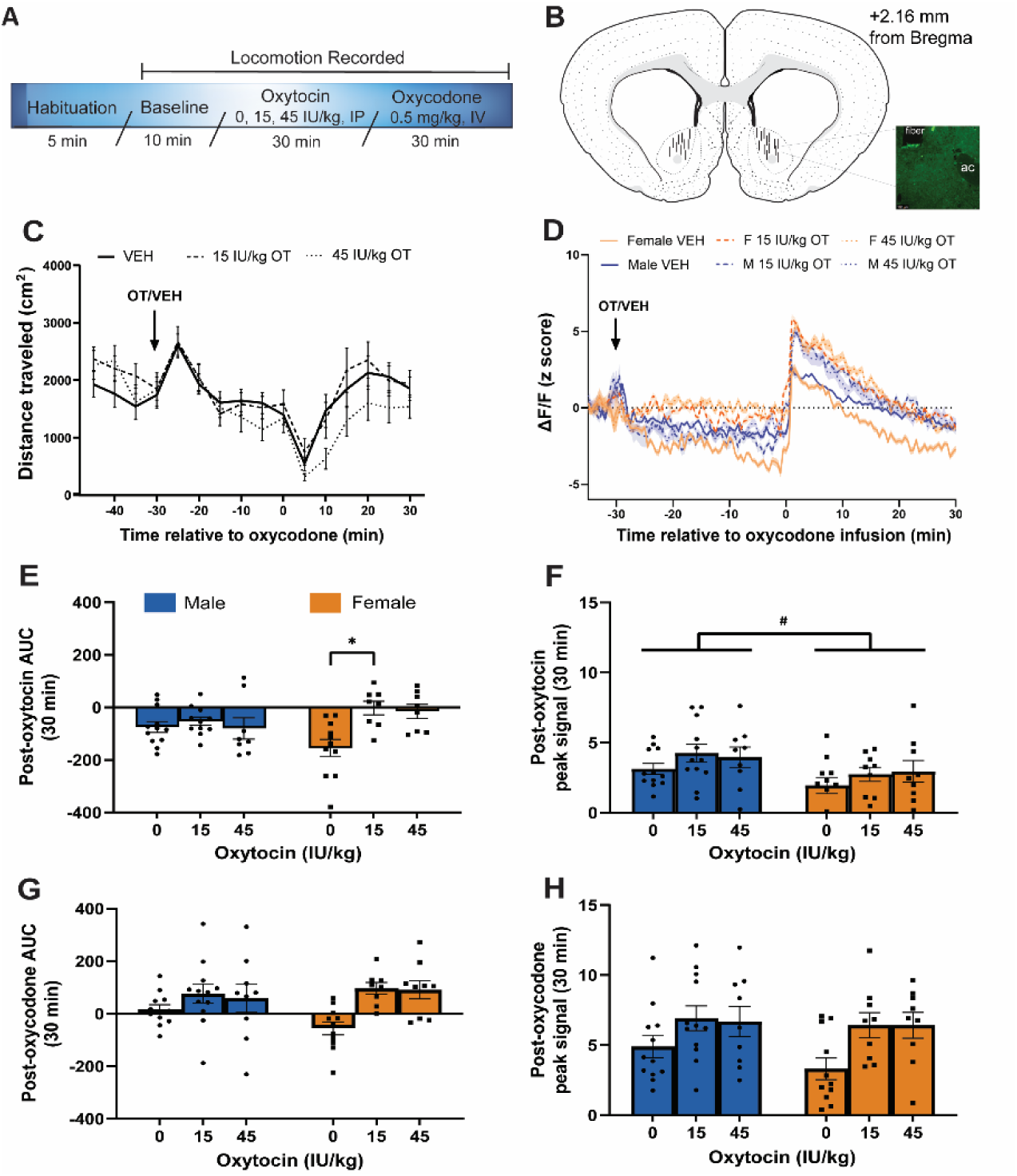
Acute oxytocin potentiated oxycodone-induced dopamine release in the nucleus accumbens core. **A.** Recording session timeline. **B.** Fiber placements in the nucleus accumbens (NA) core and a representative image of GFP-tagged GRAB_DA2m_ below the optic fiber tract, medial to the anterior commissure (ac). **C.** Neither dose of oxytocin (OT) altered locomotion compared to vehicle (VEH). **D.** Dopamine response to OT (0, 15, or 45 IU/kg) followed 30 minutes later by intravenous oxycodone. **E.** OT (15 IU/kg) increased dopamine signal area under the curve (AUC) selectively in females. **F.** Peak signal after OT/VEH was greater overall in males than in females. **G.** In both sexes, OT increased oxycodone-induced AUC (**G**) and peak signal (**H**) compared to VEH. * = p < 0.05 Female 0 IU/kg OT v. Female 15 IU/kg OT; # = p < 0.05 Female v. Male.

GRAB_DA2m_ transients were recorded using a RZ10x fiber photometry system (Tucker-Davis Technologies). Relative changes in GRAB_DA2m_ emissions (ΔF/F_0_) were extracted using custom MATLAB scripts (MathWorks) and were down sampled to 0.03 Hz as previously described (*47*). ΔF/F_0_ was then z-scored to 5 mins of baseline. Locomotion was recorded using Photobeam Activity Software (San Diego Instruments).

### Experiment 3: In vivo fiber photometry assessment of dopamine dynamics during oxycodone seeking in vehicle and acute oxytocin conditions

A separate cohort (female n = 6, male n = 9) of rats with jugular catheter implants and NA core GRAB_DA2m_ was trained to IVSA oxycodone exactly as in Experiment 1. These rats then underwent demand curve procedures to acclimate rats to high-effort responding for oxycodone reinforcement; however, unlike Experiment 1, chronic OT/VEH was not administered (see timeline in Fig. 3A). Instead, rats underwent four photometric recording sessions to determine acute OT effects on dopamine transients around lever pressing for oxycodone, using a within-subject design. Thirty min after OT (15 IU/kg, IP) or VEH, rats were tethered and placed into the operant chamber for a 1-hr recording session. OT/VEH was randomized for the first recording and then alternated between 0 and 15 IU/kg on the subsequent recordings, until rats received both doses during two tests. Oxycodone was not delivered during photometry recordings as the goal was to assess the ability of OT to reduce unreinforced lever pressing as in Experiment 1. Effort test recordings were conducted every second day, and during the interim day, rats re-acquired oxycodone IVSA on an FR-3 schedule for 3 hr/day.

**Fig. 3.**
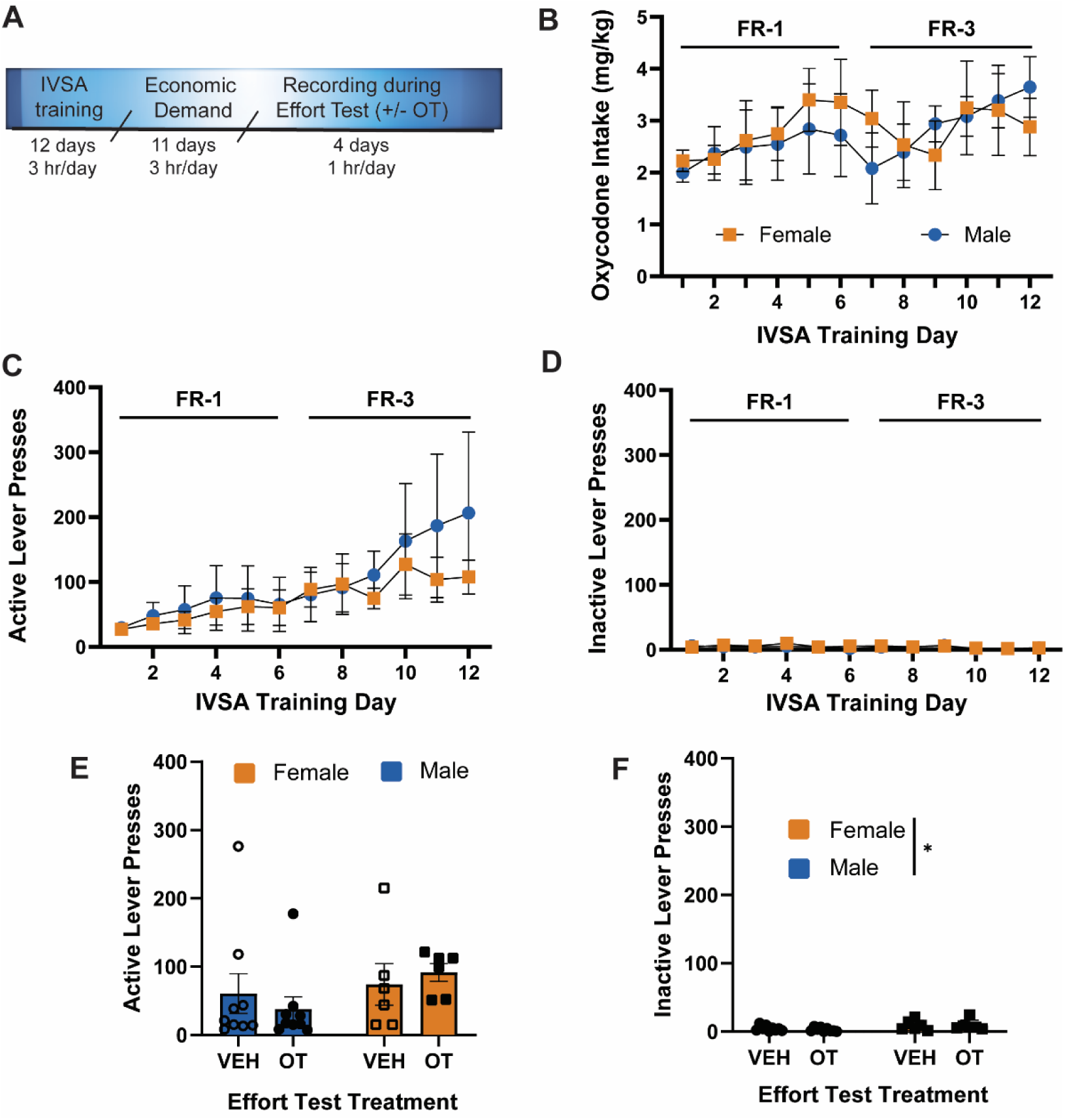
Acute oxytocin does not alter unreinforced lever pressing during fiber photometry recordings. **A.** Experimental timeline. **B.** Neither oxycodone intake, active lever presses **(C)**, or inactive lever presses **(D)** differed between females and males during intravenous self-administration (IVSA) training. **E.** 15 IU/kg oxytocin (OT) did not alter active or inactive **(F)** lever pressing between rats treated with 0 or 15 IU/kg systemic oxytocin prior to photometry recordings. Data are visualized as mean ± SEM. * = p < 0.05 Female v. Male.

Timestamps of active and inactive lever presses were transmitted to Synapse via a digital output module (DIG-726TTL, Med Associates). ΔF/F_0_ was computed and down sampled to 30 Hz. ΔF/F_0_ was z-scored to 1-second baselines, from −5 to −4 seconds before each lever press.

### Histology

Rats were administered Euthasol (100 mg/kg, IP) and were transcardially perfused with 4% paraformaldehyde. Brains were removed and 20 µm coronal sections were prepared using a Leica cryostat, mounted, and coverslipped with ProLong Diamond with DAPI. NA core cannula placement and GRAB_DA2m_ expression were verified by using a sCMOS K5 camera mounted onto a Leica DM6B epifluorescent microscope. Data from rats with fiber and AAV placement in the NA core (Figs. 2B and 4B) were included in the analysis.

**Fig. 4:**
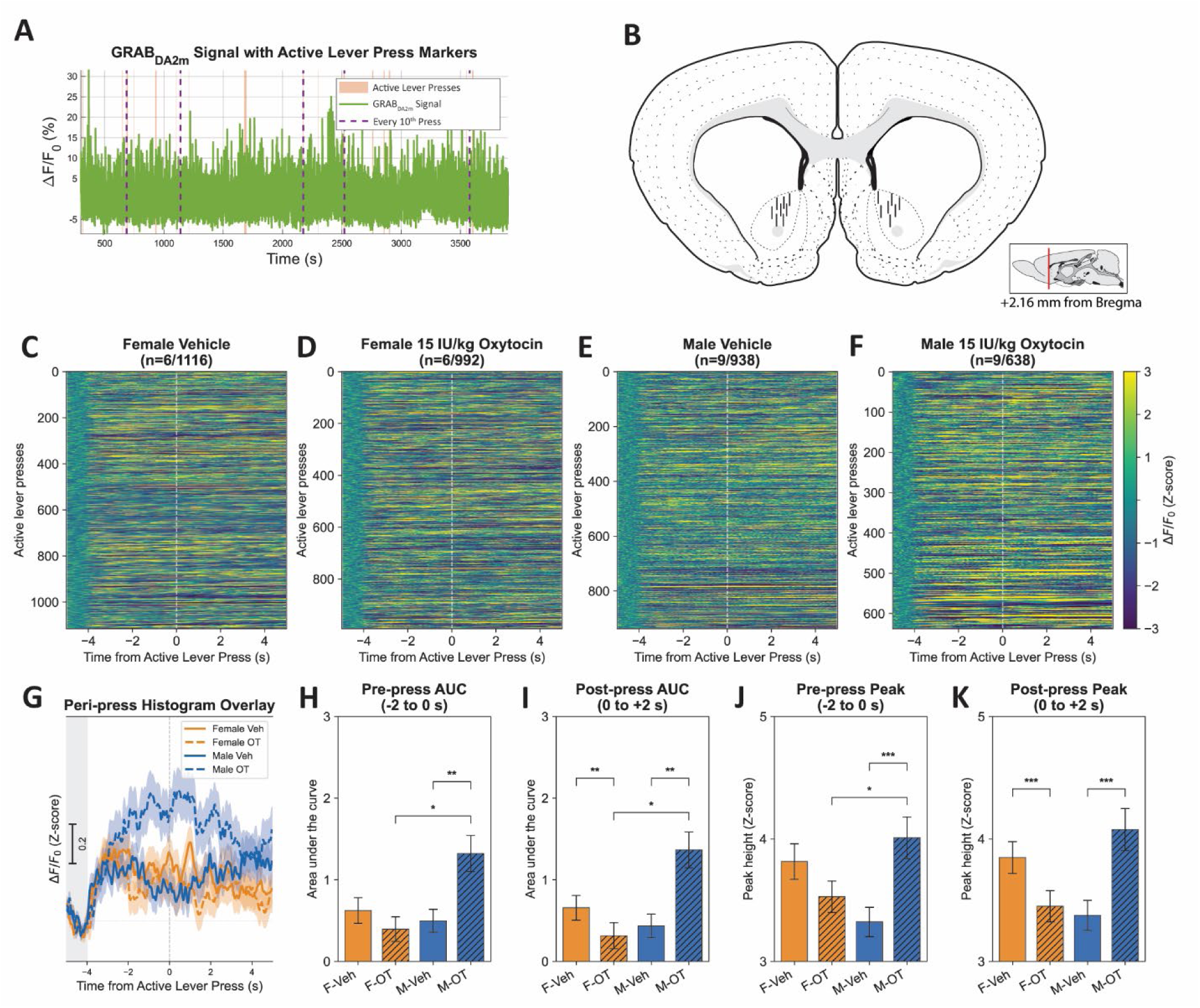
Acute oxytocin modulates peri-press dopamine dynamics during unreinforced oxycodone seeking. **A.** Representative session dopamine trace and lever press activity during a photometry recording. **B.** Fiber placements in the nucleus accumbens core. **C-F** Heatmaps illustrating dopamine dynamics around active lever press events from vehicle treated and oxytocin (OT)-treated rats. **G.** Peri-press histograms of normalized dopamine flux around active lever press events in rats treated with vehicle and oxytocin. **H.** Males treated with oxytocin (M-OT) had increased area under the curve (AUC) dopamine measures and peak signal **(J)** in the two seconds prior to active lever presses compared to vehicle-treated males (M-Veh) and oxytocin-treated females (F-OT). **I.** Males treated with oxytocin had increased area under the curve dopamine measures in the two seconds following active lever presses compared to vehicle-treated males and oxytocin-treated females. Vehicle-treated females (F-Veh) had greater area under the curve dopamine measures compared to oxytocin-treated females. **K.** Oxytocin increased male but decreased female peak dopamine measures compared to males and females treated with vehicle, respectively. * = p < 0.05 F-OT v. M-OT; ** = p < 0.01 Veh v. OT. *** = p < 0.001 VEH v. OT.

### Data analysis

GraphPad Prism (v.10.6) and SPSS (v.31) were used to conduct statistical analysis. IVSA training behavior, dopamine responses (peak, AUC), locomotion, and effort test lever pressing were compared between conditions and sex using multifactorial analyses; time was treated as a repeated measure (RM) when appropriate. Demand curve parameters (α, Q_0_ and P_max_) were quantified by fitting the exponential demand equation to infusions earned across escalating FR (k = 3.6), as previously described (*61*), and were compared between treatment groups using extra-sum of squares F tests and either independent Welch’s t- or Mann-Whitney U-tests, depending on normality of data. Significant interactions were followed with Šidák post-hoc tests controlling for multiple comparisons. Distributions were screened for outliers using ROUT (Q = 1%) and normality using Shapiro-Wilk tests. Greenhouse-Geisser corrections were used in repeated measures analyses. Level of significance was set to 0.05 and data are visualized as mean ± SEM. See Supplementary Methods for photometry analysis details.

## Results

### Experiment 1

During IVSA training, there was a main effect of Time on oxycodone intake [F_(4.467,157.2)_ = 10.02, p < 0.001; Fig. 1B] and active lever presses [F_(2.382,71.46)_ = 16.97, p < 0.001; Fig. 1C] but not inactive lever presses (Fig. 1D), indicating an escalation of intake. No effect of Sex or Sex × Time interaction was found for oxycodone intake or active and inactive lever presses during training. Rats were assigned to receive OT or VEH such that groups did not differ on any self-administration measure during training. Two male rats were eliminated due to catheter failure.

Oxytocin increased the elasticity of demand for intravenous oxycodone self-administration in males (F(_1,16_) = 59.26, p < 0.0001; Fig. 1E) but not females (F(_1,16_) = 1.792, p = 0.1994; Fig. 1F). In males, daily OT increased demand elasticity, measured by higher α [t(_14_) = 2.248, p = 0.041; Fig. 1G] and reduced P_max_ [t(_14_) = 3.551, p = 0.003; Fig. 1H] compared to VEH, indicating that OT made oxycodone seeking more sensitive to escalating price. In females, no effect of OT on α (Fig. 1I) or P_max_ (Fig. 1J) was detected. In VEH-treated rats, females showed a trend for increased α [t_(9.520)_ = 2.158, p = 0.056; Supplementary Fig. S1F] and exhibited lower P_max_ [t(_16_) = 4.031, p = 0.001; Supplementary Fig. S1G] than males. Additionally, Q_0_ of VEH-treated females was greater than male controls [U = 17, p = 0.043; Supplementary Fig. S1H].

### Experiment 2

During photometry recordings, a Sex x Dose X Time ANOVA found no effect of Sex on locomotion, and thus data is collapsed across sexes in Fig. 2C. There was no effect of Dose on locomotion. There was a main effect of Time [F_(10.50,1040)_ = 46.33, p < 0.001], with locomotion decreasing after placement into the chamber and then increasing upon IP injection, regardless of Dose, declining again, and increasing gradually after IV oxycodone administration.

Factorial mixed effects analysis of dopamine dynamics following OT/VEH and then IV oxycodone revealed a main effect of Dose [F(_2,54_)= 4.382, p = 0.017] and Time [F_(8.636, 466.3)_ = 51.18, p < 0.001] but not Sex [F(_1,54_)=0.220, p = 0.641], indicating that OT influenced dopamine dynamics in males and females in a similar manner (Fig. 2D). There were no significant two- or three-factor interactions, although several non-significant trends were detected [Time x Dose: F_(17.27, 466.3)_ = 1.506, p = 0.086; Time x Sex: F_(8.636,466.3)_ = 1.852, p = 0.060; Dose x Sex: F(_2,54_) = 2.935, p = 0.062; Time x Dose x Sex: F_(17.27,466.3)_ = 1.001, p = 0.641].

To follow up on the effect of Dose, AUC and peak signal during the 30 min following OT and OXY administration were determined. For AUC in the 30 min following OT (Fig. 2E), there was a Sex x Dose interaction [F_(1.872,29.02)_ = 4.852, p = 0.017]. OT (15 IU/kg) elevated dopamine relative to VEH in females (post hoc p = 0.015) with no significant post-hoc comparisons in males. A similar analysis of peak signal in the 30 min following OT (Fig. 2F) found a main effect of Sex (males > females) [F(_1,21_) = 5.159, p = 0.034] with no Dose effect [F_(1.741,30.46)_ = 1.919, p = 0.168] nor interaction [F_(1.741,30.46)_=0.097, p = 0.883]. As AUC *decreased* following IP injections, we then investigated the possibility that IP injections alone reduce dopamine in a separate cohort of rats (n = 3), which underwent 2 recordings each after receiving either no IP injection or IP VEH. IP injection reduced dopamine when compared to the no injection condition [Injection x Time Interaction: F_(3.341,13.37)_ = 3.628, p = 0.038; Supplemental Fig. 2A].

In the 30 min following IV oxycodone, there was a main effect of Dose [F_(1.862,31.65)_ = 11.34, p < 0.001] on AUC (Fig. 2G), with no main effect of Sex [F(_1,21_) = 0.028, p = 0.869] nor a Sex x Dose interaction [F_(1.862,31.65)_ = 2.784, p = 0.080]. Both OT doses elevated AUC relative to VEH in males and females (p’s < 0.05). Comparison of peak signal following oxycodone (Fig. 2H) found a main effect of Dose [F_(1.868,32.68)_ = 9.013, p = 0.001], with both OT doses increasing oxycodone-induced peak relative to VEH (posthoc p’s<0.05), and no effect of Sex [F(_1,21_) = 0.520, p = 0.479] nor Dose × Sex interaction [F_(1.868,32.68)_ = 0.775, p = 0.461].

### Experiment 3

Upon finding that acute OT potentiates oxycodone-induced dopamine release in both sexes, we next investigated the interaction between OT and lever pressing for oxycodone (Fig. 3A). Rats were first trained to IVSA oxycodone; during this training, male and female rats did not differ in oxycodone intake (Fig. 3B), active (Fig. 3C) or inactive lever presses (Fig. 3D). During effort testing, there was no effect of acute OT or sex on active lever pressing (Fig. 3E). There was a main effect of Sex (F > M) on inactive lever pressing [F(_1,13_) = 5.539, p = 0.035; Fig. 3F], with no main effect of OT treatment nor Sex x Treatment interaction. Thus, acute OT does not affect lever pressing for oxycodone.

Despite no effect on seeking, OT altered dopamine dynamics during unreinforced oxycodone seeking in a sex-dependent manner (Fig. 4). Active lever press associated dopamine transients from rats with placement in the NA core was analyzed (Fig. 4B). OT and sex effects on dopamine dynamics were visualized using heatmaps (Fig. C-F) and a peri-press histogram (Fig. 4G), which illustrate the most pronounced increase of dopamine signal around lever pressing in OT-treated males. AUC and peak signal during the 2 seconds prior to and following each active lever press were compared between OT conditions. Sex × Treatment interactions were found for pre-press AUC [F(_1,3038_) = 7.24, p = .007; Fig. 4H], post-press AUC [F(_1,3360_) = 18.27, p < .001; Fig. 4I], pre-press peak [F(_1,3371_) = 11.32, p < .001; Fig. 4J], and post-press peak [F(_1,3501_) = 23.93, p < .001; Fig. 4K]. Post-hoc comparisons found that OT treatment in males resulted in higher pre-press AUC, pre-press peak, post-press AUC, and post-press peak compared to VEH (p’s ≤ .001). In contrast, following OT treatment, female rats showed a decrease in post-press peak DA signal relative to the VEH condition (p = 0.047). Collectively these data suggest that OT potentiates opioid-induced NA core dopamine release equally in male and female rats, but during oxycodone seeking, OT may decrease motivation for oxycodone by selectively increasing NA core dopamine release in males.

Support vector models (SVM) were tested for the ability to predict sex and oxytocin treatment from the DA signal around lever press events (Fig. 5). SVM model performance is illustrated using receiver operating characteristic curves (Fig. 5A) and summarized in confusion matrices (Supplemental Table 1). The classifiers for vehicle treated males and females had larger AUC values than the corresponding OT classifiers in both sexes, indicating that pre-press dopamine waveform features more reliably identified active lever press events from vehicle-treated rats than from OT-treated rats. SVM classification accuracy (Supplemental Table 2) was significantly greater than scrambled data in males injected with vehicle (U = 2321, p < 0.001) and oxytocin (U = 1670, p = 0.015) but not in females, regardless of treatment (Fig. 5B). Permutation importance was calculated to identify the waveform features that contributed to SVM classification performance (Fig. 5C). The three highest-ranking features were standard deviation of amplitude, peak–valley range, and peak height. Collectively, these results indicate that amplitude-related features of NA core dopamine waveforms preceding active lever press events contributed most strongly to model discrimination between conditions.

**Fig. 5.**
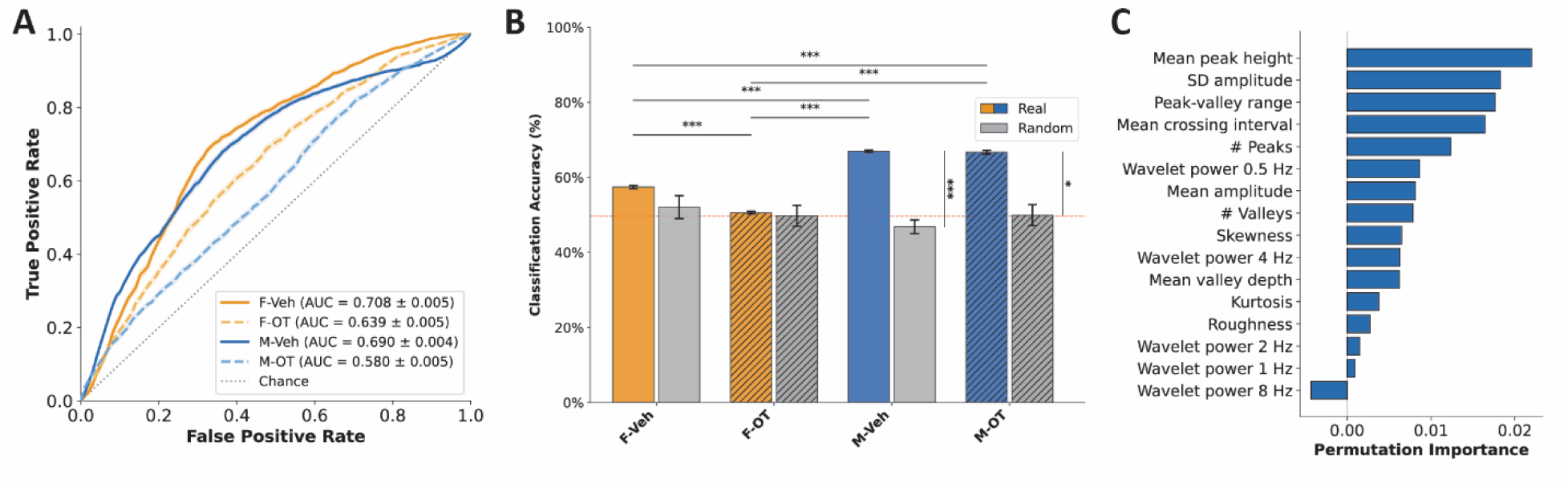
Support vector machines classify active lever presses of males in vehicle and oxytocin treatment conditions at greater than chance accuracy. **A.** Receiver operating characteristic curves of 50 support vector machines (SVM) in each treatment group illustrate superior performance in vehicle than oxytocin models. **B.** SVM accuracy is greater than scrambled data in male groups but not females. Model accuracy was greater in female oxytocin than female vehicle groups. **C.** Permutation test results of waveform features found mean peak most significantly contributed to predictive performance across models, calculated across 50 permutation repeats. Features are ranked by mean permutation importance, and larger positive values indicate greater contribution of that feature to classification performance. Data in **A-B** are visualized as mean ± SEM. Data in **C** are visualized as mean. * = p < 0.05. *** = p < 0.001.

## Discussion

Chronic oxytocin increased demand elasticity for intravenous oxycodone in males but not females. Acute oxytocin increased dopamine exclusively in females, but potentiated oxycodone-induced NA core dopamine levels in both sexes. The comparable potentiation of oxycodone-evoked dopamine flux in both sexes by acute oxytocin suggests that amplification of the dopaminergic responses to oxycodone is not, by itself, sufficient to alter oxycodone demand elasticity. During nonreinforced effort tests, acute oxytocin potentiated dopamine flux around active lever presses selectively in males, without changing lever pressing in either sex. Thus, while the behavioral effect of oxytocin on the motivation to lever press for oxycodone only emerged in males after chronic administration, the effects of acute oxytocin likely influenced the relationship between dopamine and active lever pressing, potentially influencing future oxycodone-seeking under high-effort conditions. These results implicate augmented phasic NA core dopamine signaling during drug seeking as a candidate mechanism for OUD treatment in males.

The present results supports preclinical evidence that oxytocin alters addiction-related behaviors across drug classes (*8–12*, *20*, *46*, *59*, *62–73*). Much of the existing literature has focused on acute oxytocin effects during intake, reinstatement, withdrawal, or stress-related responding, rather than the effects of prolonged oxytocin treatment on behavioral-economic factors of drug seeking and use. The present results therefore suggest that repeated oxytocin treatment can progressively reduce oxycodone seeking, specifically in males, under escalating effort requirements. The absence of an effect of chronic oxytocin in females should be interpreted with caution. Females showed more elastic oxycodone demand than males at baseline, potentially limiting any further increase in elasticity. Oxytocin receptor protein and mRNA expression is reported to be higher in males across several reward-associated regions (e.g., paraventricular nucleus of the hypothalamus, bed nucleus of the stria terminalis, lateral septum) potentially allowing for oxytocin to impact mesolimbic circuitry in a sex-specific manner (*74*, *75*). In addition, ovarian hormones regulate dopamine receptor expression and signaling, including release and transporter activity (*76–79*). Although there are some findings that suggest females are more sensitive to oxytocin than males in preclinical addiction models (*80*, *81*), other studies find no influence of sex on oxytocin effects on drug seeking/intake (*55*, *63*, *65*). While 1 and 3 mg/kg oxytocin do not reduce heroin intake in a heroin-food choice model, a sex-stratified analysis strategy was not used in that study (*20*). Accordingly, the present findings identify sex as an important variable for future oxytocin–opioid studies to consider.

Clinical trials for oxytocin in OUD have yielded mixed results. Neither acute nor chronic intranasal oxytocin decrease opioid craving in methadone or buprenorphine treated individuals with OUD (*16*, *17*). However, acute oxytocin decreases opioid craving in heroin-dependent patients (*18*), and opioid withdrawal symptoms are reduced by oxytocin (*19*). In abstinent individuals discontinuing methadone treatment, chronic oxytocin decreases opioid craving only when combined with exercise and group therapy (*15*). Thus, outcomes may depend on abstinence length, concurrent opioid agonist treatment, and psychosocial treatment context.

NA core dopamine is implicated in opioid reinforcement and seeking (*82*, *83*). Opioids increase accumbal dopamine measured by microdialysis (*50*, *84–86*), and fiber photometry. Fiber-photometry studies further show that dopamine transients track reward-predictive events and the initiation of motivated actions (*53*, *88*, *89*). Dopamine-receptor antagonism reduces opioid seeking (*37*, *38*, *87*). The present results confirm these findings, namely that acute oxycodone increases NA core dopamine, and NA core dopamine increases prior to lever pressing for oxycodone.

Acute oxytocin increased NA core dopamine around lever presses selectively in males during nonreinforced oxycodone seeking but did not acutely change responding in either sex. This dissociation may reflect a change in action-related dopamine signaling that disrupts the motivation to lever press on subsequent days. Alternatively, the OT-induced “artificial” increase in dopamine transmission around lever presses may enhance drug-reinforcement related prediction errors (*90*–*93*). Similar to the present results, VTA dopamine neuron stimulation has been reported to increase sucrose demand elasticity during within-session demand testing without reducing progressive-ratio responding (*95*).

Increased dopamine signaling can reduce reward seeking across different rewards. Systemic oxytocin enhances methylphenidate-induced dopamine release which is accompanied by attenuated methylphenidate self-administration (*96*). Drugs that increase mesolimbic dopamine like amphetamine, nicotine, and bupropion reduce sucrose self-administration (*97*, *98*). Consistent with this interpretation, pre-press dopamine transient characteristics were differentially affected by oxytocin in males and females. Together, these findings suggest that oxytocin may modify the relationship between NA core dopamine dynamics and opioid-seeking actions in a sex-dependent manner, although additional experiments are needed to determine whether these dopamine signals contribute to the chronic oxytocin-induced changes in male oxycodone demand.

In conclusion, chronic oxytocin reduced oxycodone seeking under increasing response cost in males, whereas acute oxytocin had no immediate effect on oxycodone seeking in either sex. Oxytocin potentiated oxycodone-evoked NA core dopamine signal in both sexes, increased tonic dopamine selectively in females, and enhanced phasic dopamine release around opioid-seeking selectively in males. These results support the hypothesis that chronic oxytocin may alter the relationship between NA core dopamine signaling specifically tied to lever pressing for oxycodone in a sex-dependent manner. Future work should identify the NA circuitry involved, receptor mechanisms, and whether this effect is modulated by gonadal hormones.

## Supporting information

Supplemental Materials and Results

## Data availability statement

Data will be made available upon request following publication.

## Acknowledgments

The authors thank Lizhen Wu, Alexandra Sanchez, Tyler Thompson, Talia Mandel, and Aaron Claypool for their technical assistance and Dr. Saurabh Ranjan and Dr. Micheal D. Scofield for their input on photometry analysis and coding.

## Author Contributions

HLB, CSW, MS, and LAK designed the experiments. HLB, EAC, and CSW performed surgeries and photometry. HLB and JEK conducted self-administration and histology. HLB, MPG, and CSW wrote photometry code. HLB, MPG, and LAK conducted imaging and statistical analyses. All authors contributed to writing of the manuscript.

## Funding

This work was supported by National Institute of Drug Addiction grants DA056922 awarded to LAK, DA057806 awarded to CSW, and DA059681 and DA050118 awarded to MS. The manuscript is solely the responsibility of the authors and does not represent the official views of the National Institute of Health or University of Florida.

## Competing interests

The authors declare no competing interests.

## Notes

### Competing Interest Statement

The authors have declared no competing interest.

## References

1. D. Dowell, S. Brown, S. Gyawali, J. Hoenig, J. Ko, C. Mikosz, E. Ussery, G. Baldwin, C. M. Jones, Y. Olsen, N. Tomoyasu, B. Han, W. M. Compton, N. D. Volkow, Treatment for Opioid Use Disorder: Population Estimates - United States, 2022. MMWR Morb Mortal Wkly Rep. 73, 567–574 (2024).

2. A. Manhapra, E. Agbese, D. L. Leslie, R. A. Rosenheck, Three-Year Retention in Buprenorphine Treatment for Opioid Use Disorder Among Privately Insured Adults. Psychiatr. Serv. 69, 768–776 (2018).

3. H. Wen, J. M. Hockenberry, T. F. Borders, B. G. Druss, Impact of Medicaid Expansion on Medicaid-covered Utilization of Buprenorphine for Opioid Use Disorder Treatment. Med. Care. 55, 336–341 (2017).

4. A. R. Williams, C. M. Mauro, L. Chiodo, B. Huber, A. Cruz, S. Crystal, H. Samples, M. Nowels, A. Wilson, P. D. Friedmann, R. H. Remien, M. Olfson, Buprenorphine treatment and clinical outcomes under the opioid use disorder cascade of care. Drug Alcohol Depend. 263, 112389 (2024).

5. J. Bell, J. Strang, Medication treatment of opioid use disorder. Biol. Psychiatry. 87, 82–88 (2020).

6. A. I. Mellentin, S. W. Finn, L. Skøt, D. Thaysen-Petersen, N. Mistarz, A. Fink-Jensen, D. G. Nielsen, The effectiveness of oxytocin for treating substance use disorders:A systematic review of randomized placebo-controlled trials. Neurosci. Biobehav. Rev. 151, 105185 (2023).

7. G. Gimpl, F. Fahrenholz, The oxytocin receptor system: structure, function, and regulation. Physiol. Rev. 81, 629–683 (2001).

8. M. R. Lee, C. F. Moore, E. M. Weerts, Oxytocin decreases alcohol self-administration in male baboons. Transl. Psychiatry. 14, 369 (2024).

9. C. S. Wilkinson, C. G. Modrak, T. D. Thompson, R. C. Conrad, I. Leon, L. A. Knackstedt, Consumption of oxycodone prevents oxytocin from attenuating alcohol intake in rats. Alcohol. 122, 43–53 (2025).

10. L. Zhou, W.-L. Sun, A. B. Young, K. Lee, J. F. McGinty, R. E. See, Oxytocin reduces cocaine seeking and reverses chronic cocaine-induced changes in glutamate receptor function. Int. J. Neuropsychopharmacol. 18 (2014), doi:10.1093/ijnp/pyu009.

11. G. L. Kovács, Z. Borthaiser, G. Telegdy, Oxytocin reduces intravenous heroin self-administration in heroin-tolerant rats. Life Sci. 37, 17–26 (1985).

12. D. S. Carson, J. L. Cornish, A. J. Guastella, G. E. Hunt, I. S. McGregor, Oxytocin decreases methamphetamine self-administration, methamphetamine hyperactivity, and relapse to methamphetamine-seeking behaviour in rats. Neuropharmacology. 58, 38–43 (2010).

13. M. Asker, J.-P. Krieger, A. Liles, I. C. Tinsley, T. Borner, I. Maric, S. Doebley, C. D. Furst, S. Börchers, F. Longo, Y. R. Bhat, B. C. De Jonghe, M. R. Hayes, R. P. Doyle, K. P. Skibicka, Peripherally restricted oxytocin is sufficient to reduce food intake and motivation, while CNS entry is required for locomotor and taste avoidance effects. Diabetes Obes. Metab. 25, 856–877 (2023).

14. L. Zhou, S. M. Ghee, R. E. See, C. M. Reichel, Oxytocin differentially affects sucrose taking and seeking in male and female rats. Behav. Brain Res. 283, 184–190 (2015).

15. S. Naderi, N. Vousooghi, N. Sadighii, S. A. H. Batouli, F. Mirzaii, E. Mohammad Razaghi, Oxytocin Blocks Opioid Withdrawal Symptoms Only When Combined with Group Therapy: A Double-Blind, Randomized Controlled Clinical Trial. Iran. J. Psychiatry Behav. Sci. 14 (2020), doi:10.5812/ijpbs.104627.

16. J. D. Woolley, P. A. Arcuni, C. S. Stauffer, D. Fulford, D. S. Carson, S. Batki, S. Vinogradov, The effects of intranasal oxytocin in opioid-dependent individuals and healthy control subjects: a pilot study. Psychopharmacology (Berl). 233, 2571–2580 (2016).

17. C. S. Stauffer, V. Musinipally, A. Suen, K. L. Lynch, B. Shapiro, J. D. Woolley, A two-week pilot study of intranasal oxytocin for cocaine-dependent individuals receiving methadone maintenance treatment for opioid use disorder. Addict. Res. Theory. 24, 490–498 (2016).

18. M. Moeini, A. Omidi, M. Sehat, H. R. Banafshe, The Effects of Oxytocin on Withdrawal, Craving and Stress Response in Heroin-Dependent Patients: A Randomized, Double-Blind Clinical Trial. Eur. Addict. Res. 25, 41–47 (2019).

19. B. J. Gully, Z. E. Brown, R. Hornbacher, J. C. Brown, S. E. Back, E. F. McCance-Katz, R. M. Swift, C. L. Haass-Koffler, Oxytocin Reduces Noradrenergic-Induced Opioid-Like Withdrawal Symptoms in Individuals on Opioid Agonist Therapy. Biological Psychiatry Global Open Science. 5, 100395 (2025).

20. G. Giannotti, F. Mottarlini, J. A. Heinsbroek, M. R. Mandel, M. H. James, J. Peters, Oxytocin and orexin systems bidirectionally regulate the ability of opioid cues to bias reward seeking. Transl. Psychiatry. 12, 432 (2022).

21. J. S. Carter, S. K. Wood, A. M. Kearns, J. L. Hopkins, C. M. Reichel, Paraventricular nucleus of the hypothalamus oxytocin and incubation of heroin seeking. Neuroendocrinology. 113, 1112–1126 (2023).

22. N. E. Cornejo, E. C. McNeely, T. Q. Yates, M. Kaneko, K.-C. Leong, Oxytocin attenuates yohimbine-induced responding for oral oxycodone under a progressive ratio schedule in male and female rats. Behav. Brain Res. 488, 115598 (2025).

23. D. Remillard, A. D. Kaye, H. McAnally, Oxycodone’s unparalleled addictive potential: is it time for a moratorium? Curr. Pain Headache Rep. 23, 15 (2019).

24. N. J. Connors, M. Mazer-Amirshahi, S. Motov, H. K. Kim, Relative addictive potential of opioid analgesic agents. Pain Manag. 11, 201–215 (2021).

25. B. S. Bentzley, T. C. Jhou, G. Aston-Jones, Economic demand predicts addiction-like behavior and therapeutic efficacy of oxytocin in the rat. Proc Natl Acad Sci USA. 111, 11822–11827 (2014).

26. S. B. Dolan, M. S. Berry, P. S. Johnson, M. W. Johnson, Potential for limited reinforcing and abuse-related subjective effects of intranasal oxytocin. J Psychopharmacol (Oxford). 34, 336–347 (2020).

27. R. T. Lacy, B. P. Austin, J. C. Strickland, The influence of sex and estrous cyclicity on cocaine and remifentanil demand in rats. Addict. Biol. 25, e12716 (2020).

28. J. C. Strickland, D. D. Reed, M. N. Koffarnus, S. F. Acuff, B. W. Gelino, M. E. Graham, B. A. Kaplan, A. Johansen, Z. Pierce-Messick, M. L. Banks, S. S. Negus, D. N. Kearns, D. Chiapperino, C. J. Reissig, N. B. Varshneya, S. R. Hursh, The use of behavioral economic demand in human abuse potential assessment: A review and recommendations for future testing. Exp. Clin. Psychopharmacol. (2026), doi:10.1037/pha0000877.

29. L. P. Schwartz, L. Blank, S. R. Hursh, Behavioral economic demand in opioid treatment: Predictive validity of hypothetical purchase tasks for heroin, cocaine, and benzodiazepines. Drug Alcohol Depend. 221, 108562 (2021).

30. L. R. Hammerslag, R. S. Hofford, Q. Kang, R. J. Kryscio, J. S. Beckmann, M. T. Bardo, Changes in fentanyl demand following naltrexone, morphine, and buprenorphine in male rats. Drug Alcohol Depend. 207, 107804 (2020).

31. G. F. Koob, N. D. Volkow, Neurobiology of addiction: a neurocircuitry analysis. Lancet Psychiatry. 3, 760–773 (2016).

32. C. Lüscher, R. C. Malenka, Drug-evoked synaptic plasticity in addiction: from molecular changes to circuit remodeling. Neuron. 69, 650–663 (2011).

33. L. D. Langlois, F. S. Nugent, Opiates and plasticity in the ventral tegmental area. ACS Chem. Neurosci. 8, 1830–1838 (2017).

34. J. E. Kelsey, W. A. Carlezon, W. A. Falls, Lesions of the nucleus accumbens in rats reduce opiate reward but do not alter context-specific opiate tolerance. Behav. Neurosci. 103, 1327–1334 (1989).

35. H. L. Alderson, J. A. Parkinson, T. W. Robbins, B. J. Everitt, The effects of excitotoxic lesions of the nucleus accumbens core or shell regions on intravenous heroin self-administration in rats. Psychopharmacology (Berl). 153, 455–463 (2001).

36. D. M. Hutcheson, J. A. Parkinson, T. W. Robbins, B. J. Everitt, The effects of nucleus accumbens core and shell lesions on intravenous heroin self-administration and the acquisition of drug-seeking behaviour under a second-order schedule of heroin reinforcement. Psychopharmacology (Berl). 153, 464–472 (2001).

37. J. M. Bossert, G. C. Poles, K. A. Wihbey, E. Koya, Y. Shaham, Differential effects of blockade of dopamine D1-family receptors in nucleus accumbens core or shell on reinstatement of heroin seeking induced by contextual and discrete cues. J. Neurosci. 27, 12655–12663 (2007).

38. R. T. LaLumiere, P. W. Kalivas, Glutamate release in the nucleus accumbens core is necessary for heroin seeking. J. Neurosci. 28, 3170–3177 (2008).

39. Y. Tang, Z. Chen, H. Tao, C. Li, X. Zhang, A. Tang, Y. Liu, Oxytocin activation of neurons in ventral tegmental area and interfascicular nucleus of mouse midbrain. Neuropharmacology. 77, 277–284 (2014).

40. L. Xiao, M. F. Priest, J. Nasenbeny, T. Lu, Y. Kozorovitskiy, Biased oxytocinergic modulation of midbrain dopamine systems. Neuron. 95, 368–384.e5 (2017).

41. M. R. Melis, T. Melis, C. Cocco, S. Succu, F. Sanna, G. Pillolla, A. Boi, G.-L. Ferri, A. Argiolas, Oxytocin injected into the ventral tegmental area induces penile erection and increases extracellular dopamine in the nucleus accumbens and paraventricular nucleus of the hypothalamus of male rats. Eur. J. Neurosci. 26, 1026–1035 (2007).

42. D. K. Shahrokh, T.-Y. Zhang, J. Diorio, A. Gratton, M. J. Meaney, Oxytocin-dopamine interactions mediate variations in maternal behavior in the rat. Endocrinology. 151, 2276–2286 (2010).

43. J. Peris, K. MacFadyen, J. A. Smith, A. D. de Kloet, L. Wang, E. G. Krause, Oxytocin receptors are expressed on dopamine and glutamate neurons in the mouse ventral tegmental area that project to nucleus accumbens and other mesolimbic targets. J. Comp. Neurol. 525, 1094–1108 (2017).

44. J. Rokicki, T. Kaufmann, A.-M. G. de Lange, D. van der Meer, S. Bahrami, A. M. Sartorius, U. K. Haukvik, N. E. Steen, E. Schwarz, D. J. Stein, T. Nærland, O. A. Andreassen, L. T. Westlye, D. S. Quintana, Oxytocin receptor expression patterns in the human brain across development. Neuropsychopharmacology. 47, 1550–1560 (2022).

45. C. N. Logan, G. Rojas, C. S. Wilkinson, A. K. Polo Escorcia, C. M. Reichel, J. Peris, L. A. Knackstedt, Systemic oxytocin increases glutamate efflux in the nucleus accumbens core of cocaine-experienced male and female rats but only increases dopamine efflux in males. Behav. Brain Res. 417, 113590 (2022).

46. S. T. Peters, M. T. Bowen, K. Bohrer, I. S. McGregor, I. D. Neumann, Oxytocin inhibits ethanol consumption and ethanol-induced dopamine release in the nucleus accumbens. Addict. Biol. 22, 702–711 (2017).

47. C. S. Wilkinson, S. Bhutani, W. Pyon, M. Schwendt, L. A. Knackstedt, Predator scent stress reduces oxycodone self-administration and the nucleus accumbens dopamine response to oxycodone in female rats. Addiction Neuroscience. 15 (2025), doi:10.1016/j.addicn.2025.100210.

48. F. Sun, J. Zhou, B. Dai, T. Qian, J. Zeng, X. Li, Y. Zhuo, Y. Zhang, Y. Wang, C. Qian, K. Tan, J. Feng, H. Dong, D. Lin, G. Cui, Y. Li, Next-generation GRAB sensors for monitoring dopaminergic activity in vivo. Nat. Methods. 17, 1156–1166 (2020).

49. T. Patriarchi, J. R. Cho, K. Merten, M. W. Howe, A. Marley, W.-H. Xiong, R. W. Folk, G. J. Broussard, R. Liang, M. J. Jang, H. Zhong, D. Dombeck, M. von Zastrow, A. Nimmerjahn, V. Gradinaru, J. T. Williams, L. Tian, Ultrafast neuronal imaging of dopamine dynamics with designed genetically encoded sensors. Science. 360 (2018), doi:10.1126/science.aat4422.

50. I. Willuhn, M. J. Wanat, J. J. Clark, P. E. M. Phillips, Dopamine signaling in the nucleus accumbens of animals self-administering drugs of abuse. Curr. Top. Behav. Neurosci. 3, 29–71 (2010).

51. A. A. Hamid, J. R. Pettibone, O. S. Mabrouk, V. L. Hetrick, R. Schmidt, C. M. Vander Weele, R. T. Kennedy, B. J. Aragona, J. D. Berke, Mesolimbic dopamine signals the value of work. Nat. Neurosci. 19, 117–126 (2016).

52. A. Mohebi, J. R. Pettibone, A. A. Hamid, J.-M. T. Wong, L. T. Vinson, T. Patriarchi, L. Tian, R. T. Kennedy, J. D. Berke, Dissociable dopamine dynamics for learning and motivation. Nature. 570, 65–70 (2019).

53. G. C. Touponse, M. B. Pomrenze, T. Yassine, N. Denomme, M. Wang, V. Mehta, Z. Zhang, R. C. Malenka, N. Eshel, Cholinergic modulation of dopamine release drives effortful behaviour. Nature. 651, 1020–1029 (2026).

54. C. S. Wilkinson, H. L. Blount, S. Davis, G. Rojas, L. Wu, N. P. Murphy, M. Schwendt, L. A. Knackstedt, Voluntary alcohol intake alters the motivation to seek intravenous oxycodone and neuronal activation during the reinstatement of oxycodone and sucrose seeking. Sci. Rep. 13, 19174 (2023).

55. N. A. Everett, S. J. Baracz, J. L. Cornish, The effect of chronic oxytocin treatment during abstinence from methamphetamine self-administration on incubation of craving, reinstatement, and anxiety. Neuropsychopharmacology. 45, 597–605 (2020).

56. S. Benner, Y. Aoki, T. Watanabe, N. Endo, O. Abe, M. Kuroda, H. Kuwabara, Y. Kawakubo, H. Takao, A. Kunimatsu, K. Kasai, H. Bito, M. Kakeyama, H. Yamasue, Neurochemical evidence for differential effects of acute and repeated oxytocin administration. Mol. Psychiatry. 26, 710–720 (2021).

57. J. M. Rung, M. Horta, E. M. Tammi, E. Perez, M. C. Ojeda, T. Lin, G. Harris, J. Somerville, D. Salmeron, S. E. Beltz, B. Sandesara, D. Feifel, N. C. Ebner, Safety and tolerability of chronic intranasal oxytocin in older men: results from a randomized controlled trial. Psychopharmacology (Berl). 238, 2405–2418 (2021).

58. L. W. Swanson, Brain maps 4.0-Structure of the rat brain: An open access atlas with global nervous system nomenclature ontology and flatmaps. J. Comp. Neurol. 526, 935–943 (2018).

59. K.-C. Leong, L. Zhou, S. M. Ghee, R. E. See, C. M. Reichel, Oxytocin decreases cocaine taking, cocaine seeking, and locomotor activity in female rats. Exp. Clin. Psychopharmacol. 24, 55–64 (2016).

60. L. Angioni, C. Cocco, G.-L. Ferri, A. Argiolas, M. R. Melis, F. Sanna, Involvement of nigral oxytocin in locomotor activity: A behavioral, immunohistochemical and lesion study in male rats. Horm. Behav. 83, 23–38 (2016).

61. S. R. Hursh, A. Silberberg, Economic demand and essential value. Psychol. Rev. 115, 186–198 (2008).

62. B. J. Tunstall, D. Kirson, L. J. Zallar, S. A. McConnell, J. C. M. Vendruscolo, C. P. Ho, C. S. Oleata, S. Khom, M. Manning, M. R. Lee, L. Leggio, G. F. Koob, M. Roberto, L. F. Vendruscolo, Oxytocin blocks enhanced motivation for alcohol in alcohol dependence and blocks alcohol effects on GABAergic transmission in the central amygdala. PLoS Biol. 17, e2006421 (2019).

63. N. A. Everett, A. J. Turner, P. A. Costa, S. J. Baracz, J. L. Cornish, The vagus nerve mediates the suppressing effects of peripherally administered oxytocin on methamphetamine self-administration and seeking in rats. Neuropsychopharmacology. 46, 297–304 (2021).

64. A. Bernheim, K.-C. Leong, C. Berini, C. M. Reichel, Antagonism of mGlu2/3 receptors in the nucleus accumbens prevents oxytocin from reducing cued methamphetamine seeking in male and female rats. Pharmacol. Biochem. Behav. 161, 13–21 (2017).

65. B. M. Cox, B. S. Bentzley, H. Regen-Tuero, R. E. See, C. M. Reichel, G. Aston-Jones, Oxytocin acts in nucleus accumbens to attenuate methamphetamine seeking and demand. Biol. Psychiatry. 81, 949–958 (2017).

66. A. Bahi, The oxytocin receptor impairs ethanol reward in mice. Physiol. Behav. 139, 321–327 (2015).

67. C. L. Ferland, C. M. Reichel, J. F. McGinty, Effects of oxytocin on methamphetamine-seeking exacerbated by predator odor pre-exposure in rats. Psychopharmacology (Berl). 233, 1015–1024 (2016).

68. N. A. Everett, H. A. Carey, J. L. Cornish, S. J. Baracz, Sign tracking predicts cue-induced but not drug-primed reinstatement to methamphetamine seeking in rats: Effects of oxytocin treatment. J Psychopharmacol (Oxford). 34, 1271–1279 (2020).

69. R. Ibragimov, G. L. Kovács, G. Szabó, G. Telegdy, Microinjection of oxytocin into limbic-mesolimbic brain structures disrupts heroin self-administration behavior: a receptor-mediated event? Life Sci. 41, 1265–1271 (1987).

70. S. J. Baracz, P. I. Rourke, M. C. Pardey, G. E. Hunt, I. S. McGregor, J. L. Cornish, Oxytocin directly administered into the nucleus accumbens core or subthalamic nucleus attenuates methamphetamine-induced conditioned place preference. Behav. Brain Res. 228, 185–193 (2012).

71. C. Hicks, J. L. Cornish, S. J. Baracz, A. Suraev, I. S. McGregor, Adolescent pre- treatment with oxytocin protects against adult methamphetamine-seeking behavior in female rats. Addict. Biol. 21, 304–315 (2016).

72. C. Ferrer-Pérez, A. Castro-Zavala, M. Á. Luján, J. Filarowska, R. Ballestín, J. Miñarro, O. Valverde, M. Rodríguez-Arias, Oxytocin prevents the increase of cocaine-related responses produced by social defeat. Neuropharmacology. 146, 50–64 (2019).

73. G. G. Gasanov, R. S. Ibragimov, G. L. Kovács, G. Szabó, G. Telegdy, Effect of oxytocin and its antagonist microinjected into the hippocampus on the intravenous self-administration of heroin in rats. Zh. Vyssh. Nerv. Deiat. Im. I P Pavlova. 38, 129–134 (1988).

74. J. M. Powell, K. Inoue, K. J. Wallace, A. W. Seifert, L. J. Young, A. M. Kelly, Distribution of vasopressin 1a and oxytocin receptor protein and mRNA in the basal forebrain and midbrain of the spiny mouse (Acomys cahirinus). Brain Struct. Funct. 228, 413–431 (2023).

75. C. J. W. Smith, M. L. Poehlmann, S. Li, A. M. Ratnaseelan, R. Bredewold, A. H. Veenema, Age and sex differences in oxytocin and vasopressin V1a receptor binding densities in the rat brain: focus on the social decision-making network. Brain Struct. Funct. 222, 981–1006 (2017).

76. T. M. Love, M.-A. Enoch, C. A. Hodgkinson, M. Peciña, B. Mickey, R. A. Koeppe, C. S. Stohler, D. Goldman, J.-K. Zubieta, Oxytocin gene polymorphisms influence human dopaminergic function in a sex-dependent manner. Biol. Psychiatry. 72, 198–206 (2012).

77. S. Engel, H. Klusmann, B. Ditzen, C. Knaevelsrud, S. Schumacher, Menstrual cycle-related fluctuations in oxytocin concentrations: A systematic review and meta-analysis. Front. Neuroendocrinol. 52, 144–155 (2019).

78. C. E. M. Golden, A. C. Martin, D. Kaur, A. Mah, D. H. Levy, T. Yamaguchi, A. W. Lasek, D. Lin, C. Aoki, C. M. Constantinople, Estrogen modulates reward prediction errors and reinforcement learning. Nat. Neurosci. 28, 2502–2514 (2025).

79. J. B. Becker, Gender differences in dopaminergic function in striatum and nucleus accumbens. Pharmacol. Biochem. Behav. 64, 803–812 (1999).

80. B. M. Cox, A. B. Young, R. E. See, C. M. Reichel, Sex differences in methamphetamine seeking in rats: impact of oxytocin. Psychoneuroendocrinology. 38, 2343–2353 (2013).

81. A. S. Kohtz, B. Lin, M. E. Smith, G. Aston-Jones, Attenuated cocaine-seeking after oxytocin administration in male and female rats. Psychopharmacology (Berl). 235, 2051–2063 (2018).

82. R. A. Wise, G. F. Koob, The development and maintenance of drug addiction. Neuropsychopharmacology. 39, 254–262 (2014).

83. S. A. Buck, M. M. Torregrossa, R. W. Logan, Z. Freyberg, Roles of dopamine and glutamate co-release in the nucleus accumbens in mediating the actions of drugs of abuse. FEBS J. 288, 1462–1474 (2021).

84. D. P. Devine, P. Leone, D. Pocock, R. A. Wise, Differential involvement of ventral tegmental mu, delta and kappa opioid receptors in modulation of basal mesolimbic dopamine release: in vivo microdialysis studies. J. Pharmacol. Exp. Ther. 266, 1236–1246 (1993).

85. L. Hipólito, M. J. Sánchez-Catalán, I. Zanolini, A. Polache, L. Granero, Shell/core differences in mu- and delta-opioid receptor modulation of dopamine efflux in nucleus accumbens. Neuropharmacology. 55, 183–189 (2008).

86. C. M. Vander Weele, K. A. Porter-Stransky, O. S. Mabrouk, V. Lovic, B. F. Singer, R. T. Kennedy, B. J. Aragona, Rapid dopamine transmission within the nucleus accumbens: dramatic difference between morphine and oxycodone delivery. Eur. J. Neurosci. 40, 3041–3054 (2014).

87. J. A. Heinsbroek, D. N. Neuhofer, W. C. Griffin, G. S. Siegel, A.-C. Bobadilla, Y. M. Kupchik, P. W. Kalivas, Loss of Plasticity in the D2-Accumbens Pallidal Pathway Promotes Cocaine Seeking. J. Neurosci. 37, 757–767 (2017).

88. K. M. Costa, Z. Zhang, D. Deutsch, Y. Zhuo, G. Li, Y. Li, G. Schoenbaum, Dopamine and acetylcholine correlations in the nucleus accumbens depend on behavioral task states. Curr. Biol. 35, 1400–1407.e3 (2025).

89. K. M. Fraser, V. Collins, A. R. Wolff, D. J. Ottenheimer, K. N. Bornhoft, F. Pat, B. J. Chen, P. H. Janak, B. T. Saunders, Contextual cues facilitate dynamic value encoding in the mesolimbic dopamine system. Curr. Biol. 35, 746–760.e5 (2025).

90. E. E. Steinberg, R. Keiflin, J. R. Boivin, I. B. Witten, K. Deisseroth, P. H. Janak, A causal link between prediction errors, dopamine neurons and learning. Nat. Neurosci. 16, 966–973 (2013).

91. A. S. Hart, R. B. Rutledge, P. W. Glimcher, P. E. M. Phillips, Phasic dopamine release in the rat nucleus accumbens symmetrically encodes a reward prediction error term. J. Neurosci. 34, 698–704 (2014).

92. M. P. Saddoris, F. Cacciapaglia, R. M. Wightman, R. M. Carelli, Differential dopamine release dynamics in the nucleus accumbens core and shell reveal complementary signals for error prediction and incentive motivation. J. Neurosci. 35, 11572–11582 (2015).

93. W. Schultz, Predictive reward signal of dopamine neurons. J. Neurophysiol. 80, 1–27 (1998).

94. G. L. Kovàcs, Z. Sarnyai, E. Barbarczi, G. Szabó, G. Telegdy, The role of oxytocin-dopamine interactions in cocaine-induced locomotor hyperactivity. Neuropharmacology. 29, 365–368 (1990).

95. A. M. Minnaard, M. C. M. Luijendijk, A. M. Baars, L. Drost, G. M. J. Ramakers, R. A. H. Adan, H. M. B. Lesscher, L. J. M. J. Vanderschuren, Increased elasticity of sucrose demand during hyperdopaminergic states in rats. Psychopharmacology (Berl). 239, 773–794 (2022).

96. M. R. Lee, M. C. H. Rohn, C. Zanettini, M. A. Coggiano, L. Leggio, G. Tanda, Effect of systemically administered oxytocin on dose response for methylphenidate self-administration and mesolimbic dopamine levels. Ann. N. Y. Acad. Sci. 1455, 173–184 (2019).

97. C. Fulco, C. Cordero, K. George, A. G. Roseberry, Amphetamine and nicotine reduce sucrose self-administration independent of sex. Eur. J. Pharmacol. 1020, 178796 (2026).

98. C. M. Reichel, J. D. Linkugel, R. A. Bevins, Bupropion differentially impacts acquisition of methamphetamine self-administration and sucrose-maintained behavior. Pharmacol. Biochem. Behav. 89, 463–472 (2008).

