## Supplemental Materials and Results for "Oxytocin modulates behavioral economic and dopaminergic measures of motivation to seek intravenous oxycodone in rats"

### Supplementary Information

#### Supplementary Materials and Methods

##### *Photometry data collection, processing, and analysis*

A fiber photometry system (RZ10X, Tucker-Davis Technologies, Alachua, United States) generated 415- and 465-nm GRAB<sub>DA2m</sub> sensor excitation from LED light sources using Synapse (version 100-54503) software. 415- and 465-nm light were emitted at 210 and 330 Hz frequency, respectively. Both wavelengths were set to have a 5-mA direct current offset. Light was transmitted from the light source using large bore fiber optic cables (Tucker-Davis Technologies, 200 – 600 μm core, 0.5 NA), filtered through either a LxM6 (Tucker-Davis Technologies) or FMC4 (Doric Lenses Inc., Quebec City, Canada) optical manifold, passed through a low-autofluorescence rotary joint and patch cord (Doric Lenses Inc.), and delivered to flat-tipped borosilicate fiber-optic lens (Doric Lenses Inc.) in the nucleus accumbens core. GRAB<sub>DA2m</sub> emissions were captured by the RZ10X's photodetector, digitized, and demodulated in Synapse.

Peak measures were calculated by finding the highest value in down sampled data (Experiment 2: 0.03 Hz, Experiment 3: 30 Hz), and AUC was calculated but using trapezoidal integration (e.g., MATLAB: trapz, Python: numpy.trapezoid). Active lever press time-locked GRAB<sub>DA2m</sub> signal measures were compared between treatment conditions in Python (v3.10.20) using 2-way (Sex × Treatment) linear mixed-effects models implemented through R (v4.5.3) via lme4 and a rpy2 interface. A separate cell-means model was fit for each dependent variable by restricted maximum likelihood: $DV \sim 0 + \text{Group} + (1 | \text{subject ID})$, where Group is the Sex (female, male) × Treatment (oxytocin, vehicle) factor and subject ID are a random intercept. Machine learning analyses were conducting using python, and the Mann-Whitney U test was used for comparing groupwise accuracies of support vector machine (SVM) model performance. Permutation importance and feature ranking were used to assess the contribution of each of the photometry waveform features in the SVM model, using the scikit-learn `permutation_importance` function. Permutation importance was calculated as the change in accuracy between model predictions of each feature before and after 50 random shuffles.

##### *Machine learning classification algorithm to predict biological sex and oxytocin treatment*

One-versus-rest support vector machine (SVM) classifiers were trained using pre-press dopamine waveform features to identify events originating from vehicle-treated females, OT-treated females, vehicle-treated males, or OT-treated males. Separate SVM classifiers were trained across 50 randomized train/test splits for each group to predict the drug condition (vehicle, oxytocin) and sex of rats using GRAB<sub>DA2m</sub> waveform features preceding active lever presses (-4 seconds to onset) of effort test recordings. All active lever presses were included in a clean training set if no other presses occurred within the 3 seconds before the event. Custom Python code was modified based on Kutlu et al. (2021) and primarily used the methods from Rinker et al. (2026). Various waveform features were extracted based on Maloire et al. (2019). The clean set was split into a stratified 20% held-out test set. The training set was standard scaled (fitted on the training portion, transformed on the test portion) across 50 independent models. A radial basis function-SVM ($C=1$, $\gamma = 0.001$, balanced class weight) was fitted and each model's accuracy was scored on the corresponding held-out test set. Every split was stratified uniquely and balanced to avoid bias toward the majority class. Receiving operating characteristic curves and area under the curve were used to assess discriminability of each group. These were calculated using the scikit-learn `roc_curve` and `auc` functions. Machine learning analyses were done in Python using SciPy, scikit-learn, statsmodels, and itertools.

### Supplemental Results

#### *Experiment 1*

To investigate potential sex differences in oxycodone self-administration and demand elasticity, measures were compared between male and female rats that were treated with vehicle. There was a main effect of Time on oxycodone intake [$F_{(3.971,63.54)} = 3.640$, $p = 0.010$; Supplemental Fig. 1B] and active lever presses [$F_{(2.296,36.73)} = 7.850$, $p < 0.001$; Supplemental Fig. 1C] but not inactive lever presses [$F_{(4.008,64.13)} = 1.402$, $p = 0.243$; Supplemental Fig. 1D], indicating an escalation of intake over 12 days of IVSA training. No main effect of Sex or Sex × Time interactions were found for oxycodone intake, active, and inactive lever presses during training of vehicle rats.

Vehicle treated females exhibited greater elasticity of demand for intravenous oxycodone self-administration than male controls ($F_{(1,17)} = 68.34$, $p < 0.0001$; Fig. 1E). Vehicle treated females showed a trend for increased $\alpha$ [$t_{(9.520)} = 2.158$, $p = 0.056$; Supplemental Fig. S1F] and exhibited lower $P_{\max}$ [$t_{(16)} = 4.031$, $p = 0.001$; Supplemental Fig. S1G] than males. Additionally, estimations of “free consumption”, $Q_0$ of females was greater than vehicle treated males vehicle [$U = 17$, $p = 0.043$; Supplemental Fig. S1H].

#### *Experiment 2*

Repeated measure 2-way ANOVA compared the dopamine response to IP Vehicle to a no-injection condition in a subset of 3 female rats, as illustrated in Supplemental Figure 2. An Injection × Time interaction was found [$F_{(3.341,13.37)} = 3.628$, $p = 0.038$], in addition to main effects of Time [$F_{(3.341,13.37)} = 3.536$, $p = 0.041$] and Injection [$F_{(1,4)} = 15.59$, $p = 0.017$]. Rats that received injections produced a rapid and sustained decrease in dopamine signal in the half hour following a single vehicle injection than compared to when rats were left undisturbed in the locomotor chamber.

#### *Experiment 3*

Confusion-matrix summaries of the one-versus-rest classifiers are provided in Supplemental Table 1. Across pooled held-out test observations, F-Veh and F-OT classifiers showed the highest sensitivities (79.2% and 76.1%, respectively), whereas the M-Veh classifier showed the highest precision (49.5%). The M-OT classifier had the highest specificity (74.8%) but the lowest sensitivity (34.8%), indicating that M-OT events were more often missed than correctly identified. Conversely, F-Veh and F-OT classifiers identified a high proportion of their respective positive-class observations but produced relatively high false-positive rates. Because the analysis used independent one-versus-rest models rather than a single four-class classifier, the pooled confusion-matrix summary does not identify the specific alternative group to which false-positive or false-negative events were assigned.

Supplemental Table 2 summarizes pairwise Mann–Whitney U tests comparing the distributions of held-out-set SVM accuracies across 50 randomized train/test splits, with Sidak correction applied separately to the six between-group comparisons and the four within-group real-versus-scrambled comparisons. Classification accuracy differed across most groups: F-Veh accuracy was higher than F-OT accuracy ($U = 2401.5$, Sidak-corrected $p < 0.001$) but lower than both M-Veh ($U = 13.5$, corrected $p < 0.001$) and M-OT ($U = 68.5$, corrected $p < 0.001$). F-OT accuracy was also lower than M-Veh ($U = 0$, corrected $p < 0.001$) and M-OT ($U = 1$, corrected $p < 0.001$). In contrast, classification accuracy did not differ between M-Veh and M-OT ($U = 1305$, corrected $p = 0.999$), indicating comparable performance for the two male-group classifiers. This pattern is consistent with the group means. M-Veh and M-OT achieved similar mean accuracies (0.670 and 0.667, respectively), both higher than F-Veh (0.574) and F-OT (0.506). Thus, within this waveform-feature classification framework, model performance was more strongly differentiated by sex-associated group membership than by oxytocin condition; however, this pattern should be interpreted as a difference in classifier separability rather than as a direct test of a biological sex-by-treatment effect.

Comparison of real-label and scrambled-label models further indicated that pre-press dopamine waveform features carried relevant information for male groups. Real-label classification accuracy exceeded scrambled-label performance for M-Veh ($U = 2320.5$, corrected $p < 0.001$) and M-OT ($U = 1670$, corrected $p = 0.015$), whereas F-Veh ($U = 1258.5$, corrected $p = 1.000$) and F-OT ($U = 1267$, corrected $p = 1.000$) did not differ from their scrambled-label controls. These results support above-chance classification of male vehicle- and oxytocin-treated events under the present model configuration, while evidence for reliable classification of the female groups was not found.

| Group | True Positive | False Positive | False Negative | True Negative |
| --- | --- | --- | --- | --- |
| F-Veh | 42 | 107 | 11 | 119 |
| F-OT | 51 | 124 | 16 | 89 |
| M-Veh | 54 | 55 | 38 | 133 |
| M-OT | 23 | 54 | 43 | 160 |

**Supplemental Table 1.** Confusion matrix summary results of support vector machine performance organized by group and classification outcomes.

| **Group 1** | **Group 2** | **U value** | **P value**<br>**uncorrected** | **P value Sidak-**<br>**corrected** |
| --- | --- | --- | --- | --- |
| **F-Veh** | **F-OT** | 2401.5 | 2.03E-15 | 1.22E-14 |
| **F-Veh** | **M-Veh** | 13.5 | 1.53E-17 | 9.20E-17 |
| **F-Veh** | **M-OT** | 68.5 | 3.80E-16 | 2.28E-15 |
| **F-OT** | **M-Veh** | 0 | 6.76E-18 | 4.06E-17 |
| **F-OT** | **M-OT** | 1 | 7.26E-18 | 4.35E-17 |
| **M-Veh** | **M-OT** | 1305 | 0.707 | 0.999 |
| **F-Veh** | Scrambled | 1258.5 | 0.956 | 1.000 |
| **F-OT** | Scrambled | 1267 | 0.909 | 1.000 |
| **M-Veh** | Scrambled | 2320.5 | 1.56E-13 | 6.25E-13 |
| **M-OT** | Scrambled | 1670 | 3.82E-03 | 0.015 |

**Supplemental Table 2.** Mann-Whitney U test results comparing classification accuracies between 50 support vector machine model groups.

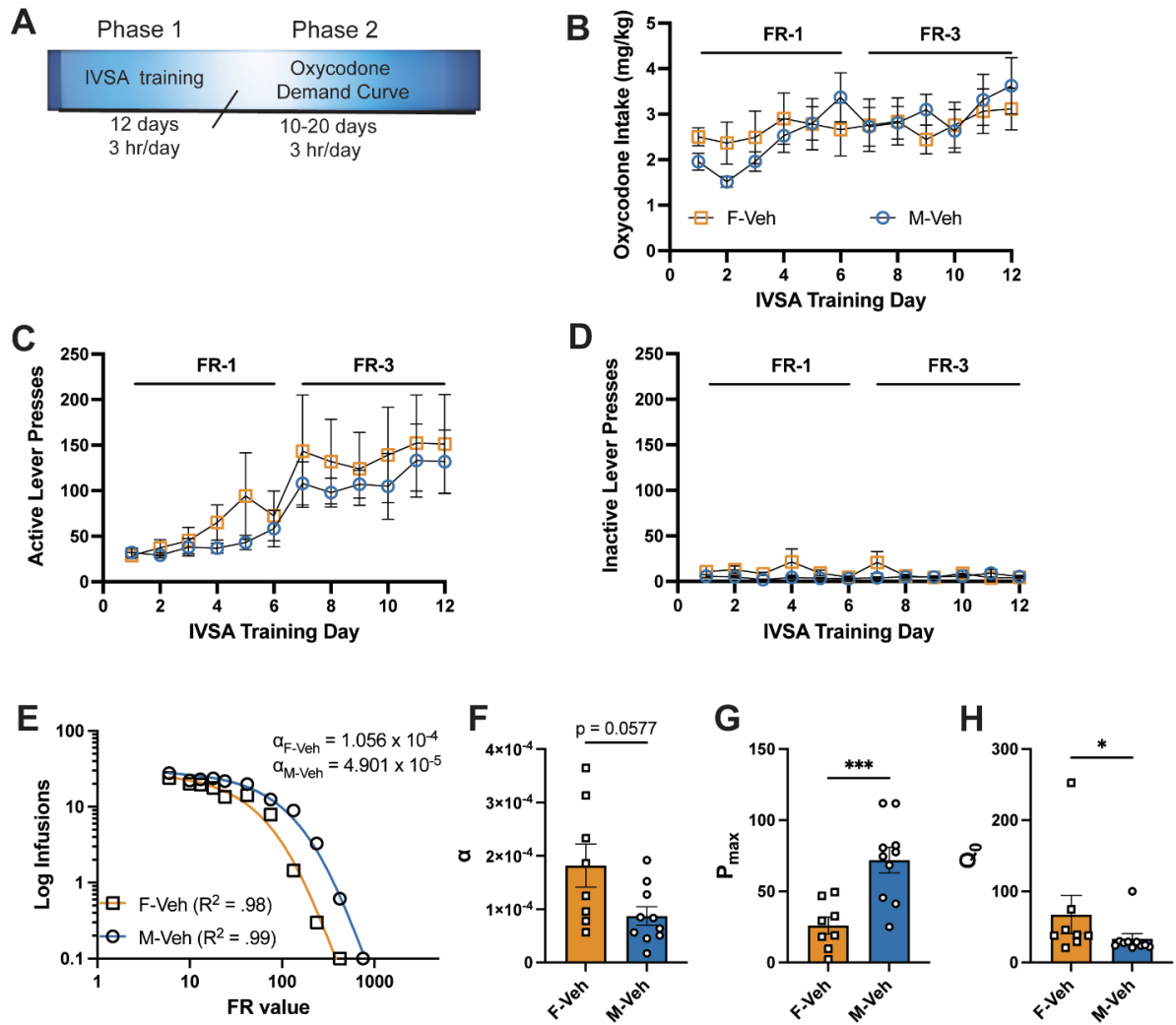

**Supplemental Figure 1.** Sex differences in economic demand for oxycodone under vehicle

conditions. **A.** Experimental timeline. During intravenous self-administration training, intake (**B**), active lever presses (**C**), and inactive lever presses (**D**) were not different between males and females later treated with vehicle in the demand curve. **E.** As oxycodone price increased between sessions of vehicle-treated controls, males were more resistant to decreasing consumption of oxycodone than females. **F.** Female vehicle-treated rats trended to have higher demand elasticity for intravenous oxycodone, had significantly lower $P_{\max}$ (**G**), and higher $Q_0$ (**H**) than males. \* = $p < 0.05$ F-Veh v. M-Veh. \*\*\* = $p < 0.001$ F-Veh v. M-Veh.

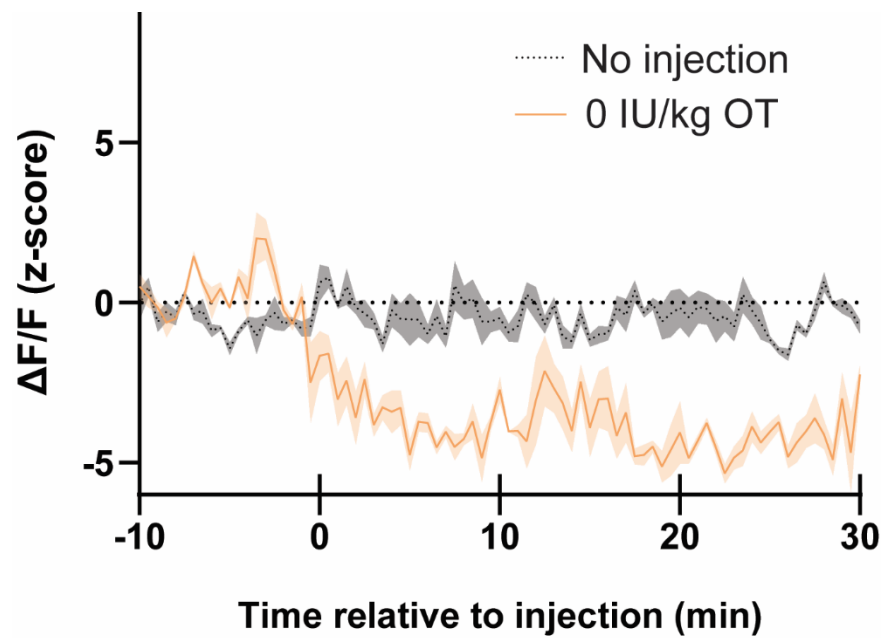

**Supplemental Figure 2.** Vehicle injections alter limbic dopamine release. Nucleus accumbens core GRAB<sub>DA2m</sub> signal of female rats decreases in the half hour following an intraperitoneal vehicle injection when compared to recordings wherein rats remained undisturbed.
